# Ligand-regulated entry into the HRD ERAD pathway—the dark side of allostery

**DOI:** 10.1101/177311

**Authors:** Margaret A Wangeline, Randolph Y Hampton

## Abstract

HMG-CoA reductase (HMGR) undergoes regulated degradation as part of feedback control of the sterol pathway. In yeast the stability of the Hmg2 isozyme of HMGR is controlled by the 20 carbon isoprenoid geranylgeranyl pyrophosphate (GGPP): increasing levels of GGPP causes more efficient degradation by the HRD pathway, allowing feedback regulation of HMGR. The HRD pathway is a conserved quality control pathway critical for the ER-associated degradation of misfolded ER proteins. We have explored the action of GGPP in HRD-dependent Hmg2 degradation. GGPP was highly potent as a regulatory molecule in vivo, and *in vitro*, GGPP altered Hmg2 folding at nanomolar concentrations. These effects of GGPP were absent in a variety of stabilized or non-regulated Hmg2 mutants. Consistent with its high potency, the effects of GGPP were highly specific; other closely related molecules were ineffective in altering Hmg2 structure. In fact, two close GGPP analogues, 2F-GGPP and GGSPP were completely inactive at all concentrations tested, and GGSPP was an antagonist of GGPPs effects *in vivo* and *in vitro*. The effects of GGPP on Hmg2 structure and degradation were reversed by chemical chaperones, indicating that GGPP caused selective Hmg2 misfolding. These data indicate that GGPP functions in a manner analogous to an allosteric ligand, causing Hmg2 misfolding through interaction with a reversible, specific binding site. Consistent with this, the Hmg2 protein forms mulitmers. We propose that this “allosteric misfolding,” or *mallostery*, may be a widely used tactic of biological regulation, with potential for development of small molecule pharmaceuticals that induce selective misfolding.

## INTRODUCTION

Protein quality control includes a variety of mechanisms to ensure tolerably low levels of misfolded proteins in the living cell. Among these, selective degradation of misfolded, damaged or un-partnered proteins is often employed for removal of these potentially toxic species. One of the best characterized pathways of degradative quality control is ER-associated degradation (ERAD), entailing a group of ubiquitin-mediated pathways that degrade both lumenal and integral membrane proteins of the endoplasmic reticulum (ER) (1–4). All degradative quality control pathways show a remarkable juxtaposition in their action. They are all highly specific for misfolded versions of the substrate proteins, yet they recognize a wide variety of distinct and unrelated substrates (5, 6). This “broad selectivity” is based on the ability of the ubiquitination enzymes to recognize or respond to specific structural hallmarks of misfolding shared by a wide variety of client substrates. The details and restrictions of these recognition features are still being discovered due to the apparently wide range of ways that E3 ligases can detect their clients (5, 7–9).

The remarkable selectivity for misfolded proteins positions degradative quality control as a powerful tactic for physiological control of normal proteins. It is now clear that a number of cases exist where a normal protein can enter a bone fide quality control pathway to bring about its physiological regulation (10–16). The best studied example of this sort of control is the regulated degradation of HMG-CoA reductase (HMGR), a rate limiting enzyme of the sterol synthetic pathway. In both mammals and yeast, this essential enzyme undergoes regulated degradation in response to molecular signals from the sterol pathway as a mode of feedback control of sterol synthesis. In both cases, ERAD pathways are employed to bring about the regulated degradation of the normal enzyme, allowing for a deep understanding of selectivity in ERAD, and holding the promise for development of new strategies to control the levels of individual protein targets.

Our studies of sterol regulation in *S. cerevisiae* show that the HRD ERAD pathway mediates the regulated degradation of the Hmg2 isozyme of HMGR. The HRD pathway is centrally involved in mitigating ER stress through ubiquitin-mediated degradation of a wide variety of misfolded, ER resident lumenal and integral membrane proteins (12, 17–19). The primary signal for Hmg2 degradation is the 20 carbon sterol pathway product geranylgeranyl pyrophosphate (GGPP) (Fig 1A) which is produced during normal cell anabolism and is thus a fiduciary indicator of sterol pathway activity (20). When levels of GGPP are high, HRD dependent degradation of Hmg2 increases, and when GGPP levels are low, Hmg2 becomes more stable, thus effecting feedback control at the level of enzyme stability. It was initially surprising that the broadly used HRD quality control pathway is required for the precisely regulated degradation of normal Hmg2. Because the HRD pathway functions to remove misfolding proteins, we had previously posited that GGPP functions by promoting a change in the structure of Hmg2 to a better HRD pathway substrate, thus employing the selectivity of the HRD machinery for purposes of physiological regulation. The studies herein test and explore that idea.

**Figure 1.**
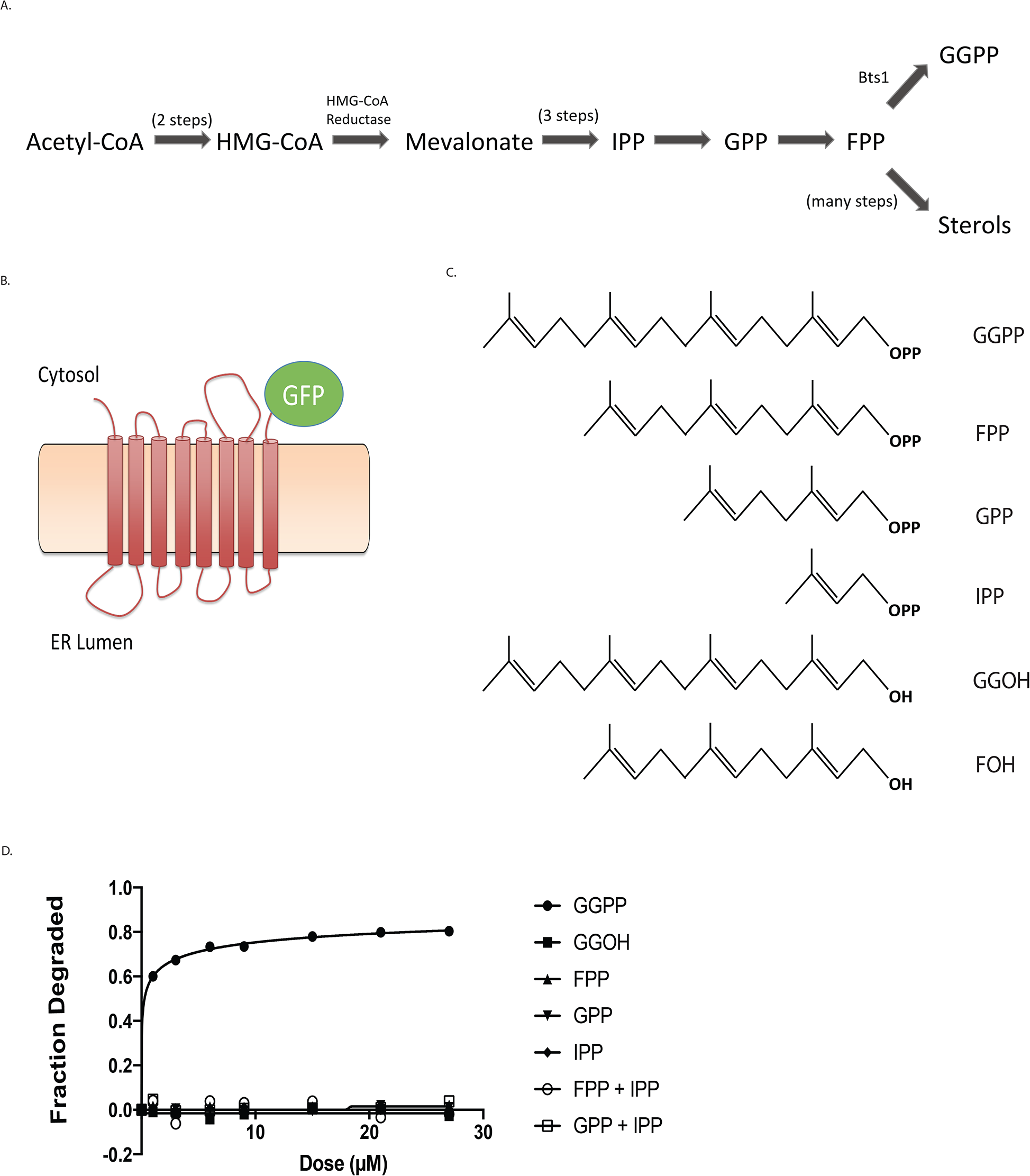
Effect of GGPP and related molecules on *in vivo* Hmg2-GFP degradation. (A) Schematic of the basic sterol biosynthetic pathway, showing relative positions of GGPP and other molecules mentioned in the studies. (B) Representation of Hmg2-GFP, a multispanning integral membrane ER protein and HRD pathway substrate. Hmg2-GFP undergoes normal regulated degradation, but has no catalytic activity. In this construct, GFP replaces the C-terminal cytoplasmic catalytic domain. (C) Structures of key isoprenoid molecules studied in this work, including the potent biological regulator of Hmg2, GGPP. (D) Dose-response curve of the molecules pictured in 1c for stimulateing Hmg2-GFP degradation, indicated by loss of *in vivo* fluorescence after a 1 hour 30° C incubation after direct addition of indicated compounds to culture medium followed by flow cytometry, counting 10,000 cells. Fractional degradation is the difference in fluorescence initially versus 1 hour, as a fraction of initial fluorescence.

**Figure 2.**
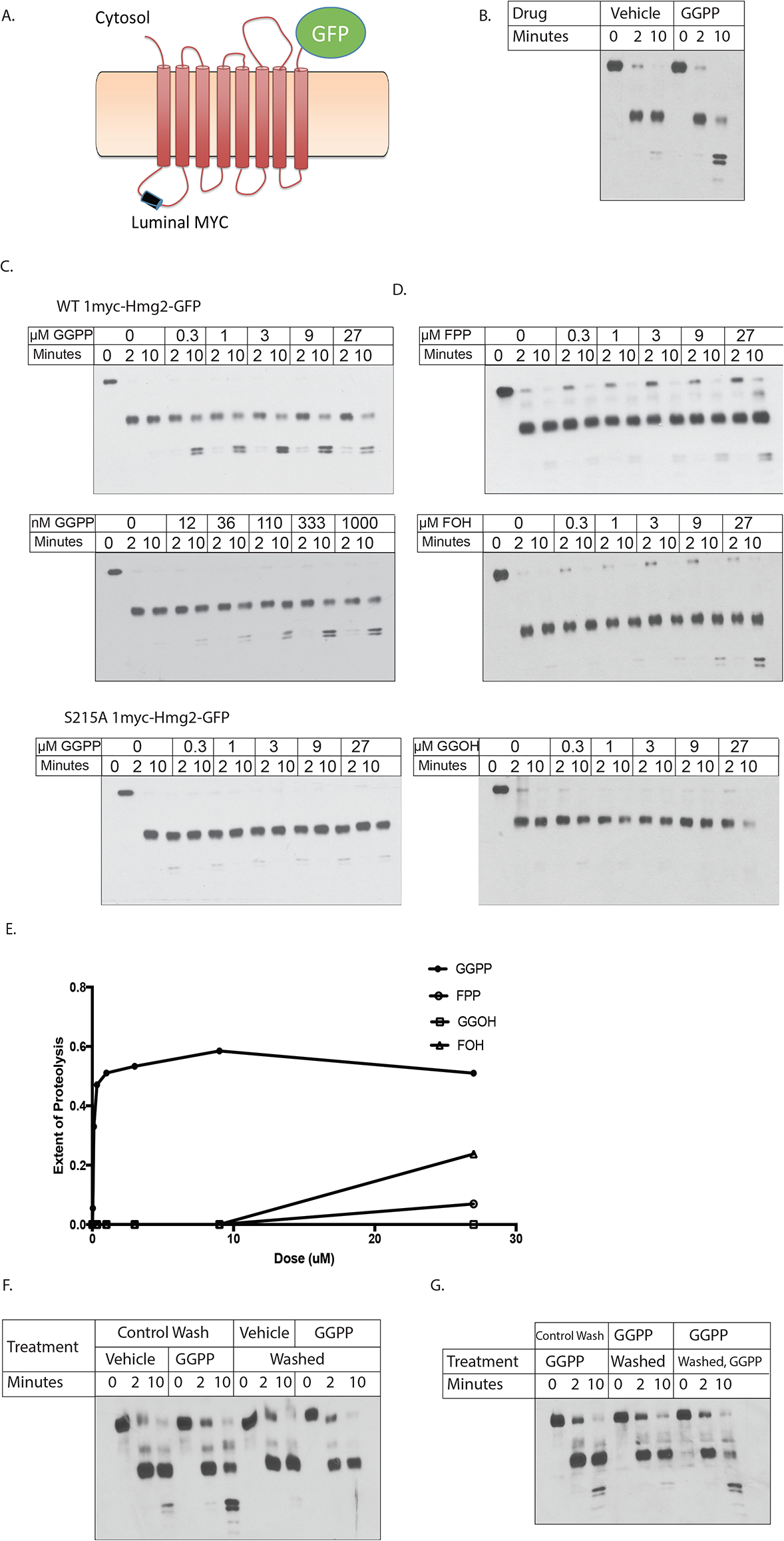
Effect of GGPP and related molecules on Hmg2-GFP structure *in vitro*. (A) Cartoon of 1myc-Hmg2-GFP. Fully regulated Hmg2-GFP with protected lumenal 1myc epitope to allow limited proteolysis assay of Hmg2 structure in isolated microsomes, as described below and in (25). (B) Effect of GGPP on limited proteolysis assay. 9 μM GGPP added to microsomes immediately before incubation with trypsin, followed by SDS-PAGE and myc immunoblotting as described. (C) Concentration dependence of of GGPP on wild-type 1myc-Hmg2-GFP with normal transmembrane region or the highly stabile S215A point mutation in the SSD. D) Effect of some other molecules on Hmg2-GFP limited proteolysis (D), In all panels, molecule being tested and concentration range employed is listed in upper left corner; concentrations are indicated along panel top. Note effect of GGPP begins at approximately12 nM. (E) Graphical representation of results in 2 C,D. Immunoblot films were scanned as TIFF files, and pixel counted for total lane intensity and pixels in final doublets that result from extended incubation with trypsin. Extent of proteolysis = (doublet intensity/total lane intensity). (F,G) GGPP effect on Hmg2-GFP structure is fully reversible, and GGPP responsiveness remains after reversal. 2F: Microsomes were washed and then treated with vehicle or GGPP (groups 1 and 2), or treated with vehicle or GGPP and then washed (groups 3 and 4). All groups were then subjected to limited proteolysis assay as described. Note that washing first did not affect response to GGPP, while washing after exposure removed effect. 2G: Microsomes that were treated with GGPP and washed maintained their ability to respond to readdition of GGPP. Left set of microsomes was washed, then treated with GGPP (left group). Middle set was treated with GGPP then washed, and the right group was treated with GGPP, washed, then re-treated with GGPP. All samples were then subjected to limited proteolysis assay. Note re-addition of GGPP gave precisely the same response as first addition.

We found that indeed GGPP directly influenced the structure of the Hmg2 multispanning anchor, in the low-to-mid nanomolar range. These potent actions of GGPP were highly specific, and in fact were antagonized by a close GGPP analogue both *in vivo* and *in vitro*. Furthermore, the effects of GGPP were blocked by a variety of chemical chaperones, indicating that this molecule causes remediable misfolding of the Hmg2 structure to promote HRD recognition. Taken together, these studies lead to a natural model of regulated quality control as a form of allostery that may be widely employed in biology to harness the intrinsic specificity of the many branches of degradative quality control. Because this axis of regulation appears to be based on reversible misfolding due to specific ligand binding, we have given it the name “mallostery” to reflect both the elements of misfolding implied by the prefix, and the action of a selective regulatory ligand that hallmarks allosteric control of many enzymes and other proteins.

## RESULTS

*Specificity and potency of isoprenoids that stimulate Hmg2 degradation-In* our earlier work, we tested the effects of a variety of sterol pathway molecules on Hmg2 stability (20, 21). We found that only the 20-carbon isoprenoid geranylgeranyl pyrophosphate (GGPP) caused Hmg2 degradation *in vivo* when added to culture medium (Garza *et al*, 2009). This surprising ability of exogenous GGPP to stimulate Hmg2 degradation has been a useful feature for study of this regulatory signal (22, 23). Because this response is part of a selective negative feedback loop, we posited that the GGPP signal would be specific, physiologically relevant, and highly potent. To more systematically evaluate these ideas, we first performed dose response experiments on pathway isoprenoids alone and in combination.

We examined the effects of candidate isoprenoids on Hmg2 stability *in vivo* using flow cytometry on cells expressing Hmg2-GFP, which undergoes regulated degradation identical to the native enzyme (24), but provides no additional enzymatic contribution to signal production. Each was tested at a variety of concentrations by direct addition to yeast cultures, followed by a 1 hour incubation and flow cytometry. GGPP caused Hmg2-GFP degradation at culture concentrations as low as 1 μM, reaching a maximum at approximately 20 μM (Fig 1C). The effect of GGPP on Hmg2-GFP was highly specific: the 15 carbon farnesyl pyrophosphate (FPP) and the non-phosphorylated 20 carbon geranylgeraniol (GGOH) had no effect *in vivo*. Similarly, neither of the earlier pathway isoprenoids, isopentenyl pyrophosphate (IPP) or geranyl pyrophosphate (GPP), had any effect on Hmg2-GFP at concentrations up to 27 μM.

Because GGPP is synthesized by addition of the 5 carbon IPP (Fig 1A) to FPP, we also tested if addition of both FPP and either of the interconvertible precursors might simulate direct addition of GGPP by allowing synthesis of this regulator from these precursors. Accordingly, we also treated cells simultaneously with the combinations IPP and FPP, or GPP and FPP. Neither of these co-additions had any effect.

These results indicated a clear structure-function relationship for GGPP as a degradation signal, since similar molecules did not act to stimulate Hmg2 degradation. We were curious how stringent the structural features of GGPP were, so we next tested two close analogues of GGPP: 2F-GGPP and GGSPP (Fig 3A). Despite the striking similarity to GGPP, neither of these molecules stimulated Hmg2 degradation *in vivo* at even very high concentrations. Thus, the *in vivo* effect of GGPP on Hmg2 degradation appeared to be highly specific. The high specificity of GGPP the *in vivo* assay could have a variety of explanations so we turned to our previously employed *in vitro* assay to directly evaluate the action of GGPP on regulated stability of Hmg2.

**Figure 3.**
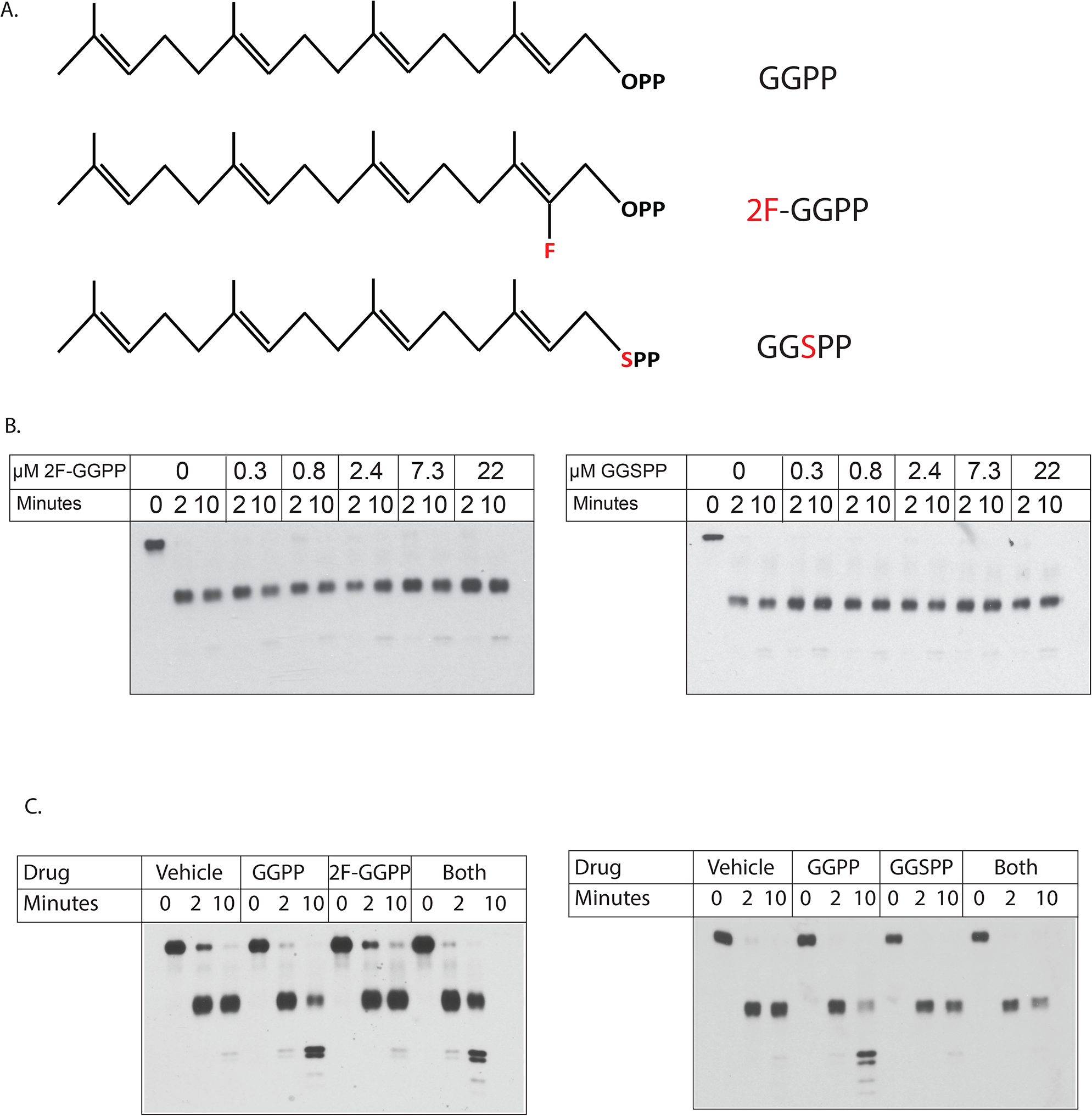
Inactive and antagonistic analogues of GGPP in Hmg2 structural assay. (A) Structures of GGPP, 2F-GGPP and GGSPP, tested for activity and antagonism in this figure; note identical 20 carbon lipid groups of each. (B) Dose-response of the two inactive GGPP analogues in limited proteolysis assay. Note neither compound had any effect even at the highest concentration tested (22 μM). (C) Test of each inactive GGPP analogue for antagonism of GGPP effect in limited proteolysis assay. On both panels, 3 μM GGPP was added where indicated, along with 44 M of indicated analogue. Microsomes were tested for effects of no addition, addition of each molecule, or the simulateneous addition of GGPP and an analogue. 2F-GGPP was tested on the left panel, and GGSPP was tested on the right panel. Note only GGSPP antagonized the effect of GGPP.

*In vitro analysis of GGPP action on Hmg2-Our* early studies described a limited proteolysis assay for studying the effects of small molecules and expressed proteins on the structure of the Hmg2 transmembrane domain (25, 26). The assay uses myc_L_-Hmg2-GFP, a version of Hmg2-GFP with an added single myc tag inserted into the first luminal loop of the transmembrane domain (Fig 2A). The exact placement of lumenal tag along the Hmg2 sequence provides two key features: first, it does not perturb *in vivo* regulation of the resulting protein. Second, because the myc tag is present in the lumenal space, complete proteolysis of the tagged Hmg2 can be accomplished by addition of proteases to the cytoplasmic side of ER-derived microsomes without loss of myc signal (25). Because ER microsomes from yeast are almost completely cytosol-side-out (27), expression of myc_L_-Hmg2-GFP allows facile analysis of structural features of microsomal Hmg2-GFP with a simple limited proteolysis assay (22, 25, 26, 28).

When microsomes isolated from cells expressing myc_L_-Hmg2-GFP are treated with a low concentration of trypsin, immunoblotting the protected myc epitope after SDS-PAGE reveals a characteristic time-dependent pattern of proteolyzed fragment production (Fig 2B). Because the myc tag is protected, the total myc immunoblotting signal intensity remains unchanged. We developed this assay to explore how signals from the sterol pathway affect the structure of Hmg2 to render it more susceptible to the HRD quality control pathway (26). In those early studies we found that the rate of myc_L_-Hmg2-GFP proteolysis was altered by manipulations that affect the *in vivo* stability of the protein, such that *in vitro* proteolysis occurred more rapidly when microsomes were prepared from strains where the degradation signals are high (26). *In vivo*, Hmg2 or Hmg2-GFP is strongly stabilized by chemical chaperones (29). Similarly, proteolysis of microsomal myc_L_-Hmg2-GFP is drastically slowed by addition of the chemical chaperone glycerol, and this structural change is fully reversible (25). We employed this *in vitro* structural assay to explore the possibility that sterol pathway signals directly affected the structure of Hmg2 to allow regulated degradation.

In those studies, we showed that the 15 carbon neutral isoprenoid farnesol (FOH) caused significant acceleration of *in vitro* myc_L_-Hmg2-GFP trypsinolysis, again preserving the cleavage pattern but altering the kinetics (26). This effect of FOH is fully reversible. Furthermore, mutants of Hmg2-GFP that do not respond to *in vivo* degradation signals, including a substitution of a small number of amino acids known as “TYFSA”, or a single point S215A point mutant of a highly conserved residue of the sterol-sensing domain (SSD), do not respond to farnesol in the limited proteolysis assay (22, 26). Although those results were intriguing and biologically appropriate, the biological role of farnesol per se was unclear. Although there was a clear structure-activity relationship for farnesol in the proteolysis assay, the concentrations required to cause the *in vitro* effects were very high (EC_50_ ~ 100 uM), and farnesol is extremely toxic to yeast. In the times since these studies, we discovered that the *bona fide* physiological regulator was the normally made isoprenoid GGPP, which also causes the structural transition of Hmg2-GFP in the proteolysis assay (20). Accordingly we returned to this assay to evaluate the specificity and potency of GGPP in a more controlled setting.

In striking contrast to FOH, we found that GGPP was a potent modifier of Hmg2 structure. GGPP accelerated *in vitro* trypsinolysis at concentrations as low as ~15 nM, with an apparent half-maximum concentration lower than 200 nM (Fig 2C left, 2E). Intriguingly, this concentration is in the range of the K_M_ of yeast enzymes that use GGPP as a substrate (30–32), indicating that this concentration is likely physiologically relevant since the enzymes are “tuned” to concentrations of substrate that exist in their milieu. The maximal effect of GGPP was similar to that seen with the largest effects of FOH reported earlier, about a 5 fold increase in proteolysis rate. The highly stable mutant S215A, which does not respond to FOH in the *in vitro* assay, also did not respond to GGPP at any concentration tested (Fig 2C, right).

To investigate the specificity of this potent effect of GGPP, we directly compared a variety of isoprenoid molecules, GGOH, FPP, and FOH all of which we have previously shown accelerate *in vitro* trypsinolysis to some degree. While all three, to varying degrees, altered Hmg2-GFP structure, their effects occurred at half-maximum concentrations over 100 fold higher than GGPP (Fig 2D,E). We next tested the close structural analogues of GGPP, 2-fluoro GGPP (2F-GGPP) and S-thiolo GGPP (GGSPP), since these were inactive in the *in vivo* assay. Even in the direct *in vitro* assay, neither analogue induced the structural transition at any concentration tested, over 40 μM (Fig 3B,C).

The much higher potency of GGPP compared to isoprenoid alcohols and non-hydrolyzable analogs raised the question of whether GGPP was being used to covalently modify Hmg2 or another protein in the microsome extract. The yeast geranylgeranyl transferase machinery is cytosolic, and thus unlikely to have activity in our washed membranes. Nevertheless, we addressed this possibility by testing whether the GGPP effect on Hmg2-GFP was readily reversible. Microsomes were prepared normally and then washed three times in reaction buffer either before or after treatment with vehicle and GGPP. When microsomes were washed before treatment with GGPP, Hmg2-GFP became more susceptible to proteolysis as usual. When microsomes were washed after treatment, GGPP’s effect was reversed (Fig 2F). Furthermore, the treated and then washed microsomes remained competent for the GGPP-induced structural transition: when GGPP was added back to the washed microsomes, Hmg2-GFP again became more susceptible to proteolysis, and to the same extent as the original exposure (Fig 2G).

*Antagonism of GGPP action in vitro and in vivo-*The GGPP analogues 2F-GGPP and GGSPP had no ability to stimulate Hmg2-GFP degradation *in vivo* (Fig 4A,C), nor alter Hmg2-GFP structure in the limited proteoloysis assay (Fig 3B). The high potency and specificity of GGPP, and its ability to directly and reversibly alter the structure of Hmg2-GFP made us wonder if it acts as a ligand, causing a structural change by specific interaction with the Hmg2 transmembrane region at a particular binding site, similar to allosteric regulation of enzymes by regulatory metabolites. Accordingly, we asked if an excess of either of the highly similar, inactive analogues might antagonize the effects of GGPP. Each was tested for an ability to block the structural effect of a low concentration of GGPP by co-incubation with an excess of analog. As expected, the test doses of either analog had no effect on mycL-Hmg2-GFP (Fig 3B). However, the presence of a 15-fold molar excess of GGSPP clearly antagonized the structural effect of GGPP. Interestingly, only one of the analogues had this effect; the 2F-GGPP was simply inactive in an identical experiment (Fig 3C). This is particularly important since both molecules have very similar chemistry and amphipathicity, both being developed to block the same class of enzymes (33, 34). Nevertheless, only GGSPP antagonized the GGPP-induced structural effects on mycL-Hmg2-GFP.

**Figure 4.**
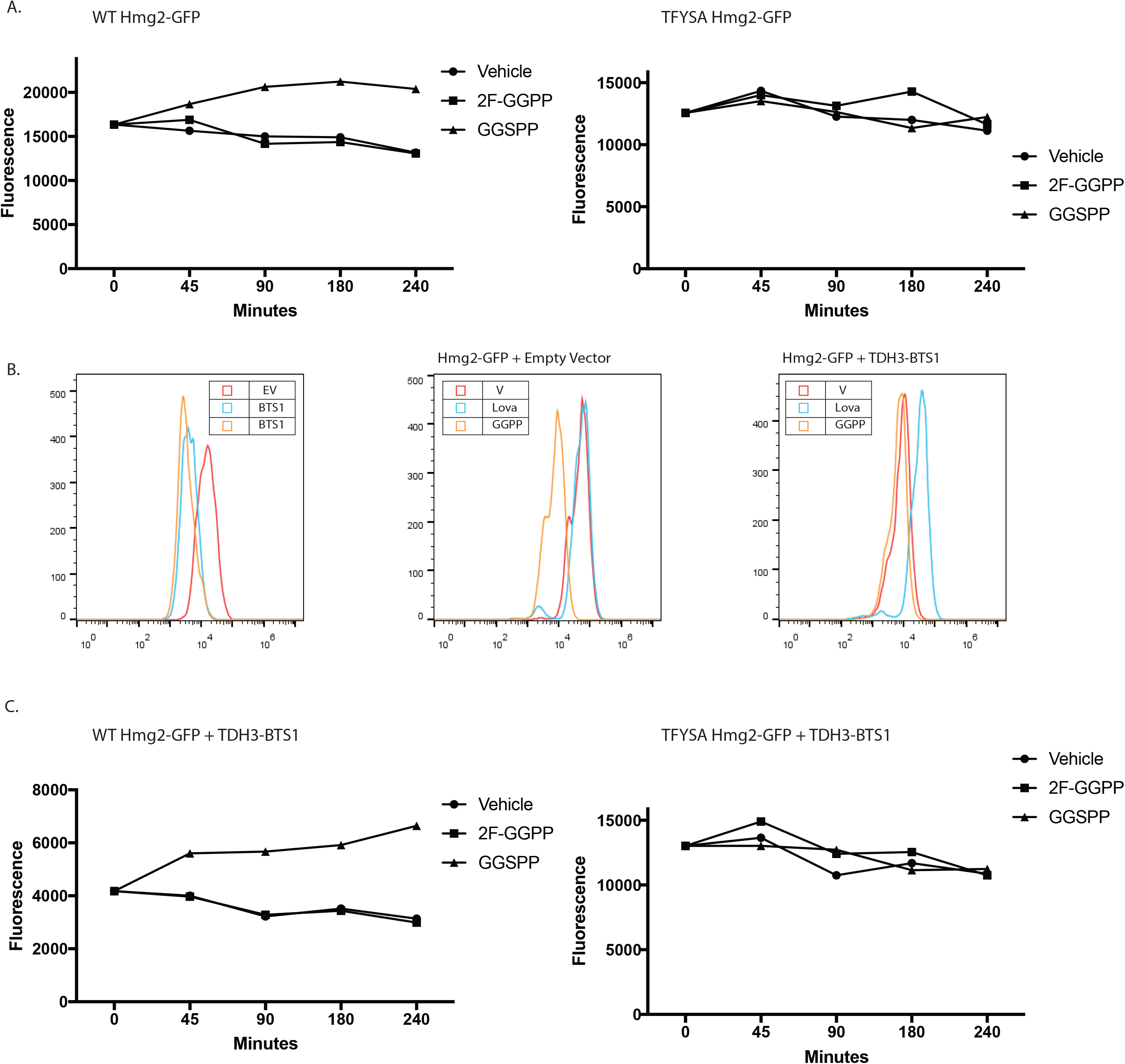
*In vivo* antagonism of GGPP-stimulated Hmg2-GFP degradation. (A) Effect of 2F-GGPP (non-antagonist *in vitro)* or GGSPP (antagonist *in vitro*) on wild type strain with normal production of GGPP due to the sterol pathway, expressing regulated Hmg2-GFP (left panel) or non-regulated TYFSA mutant of Hmg2-GFP (right panel). All graphs show mean fluorescence by flow cytometry of 10,000 cells. (B) Strains with elevated GGPP production due to strongly expressed BTS1 gene encoding GGPP synthase. Left panel, steady-state fluorescence of strains expressing empty vector (red) or integrated BTS 1 expression plasmid (blue, orange), showing strong shift in steady-state level of Hmg2-GFP fluorescence due to elevated endogenous GGPP production. Middle panel, effect of lovastatin (blue) or GGPP (orange) on Hmg2-GFP fluorescence on empty vector strain. Right panel, same experiment with a BTS1-expressing plasmid present; note addition of GGPP has little further effect and that lovastatin, that blocks GGPP production due to elevated BTS 1, causes strong stabilization. (C) Effect of GGPP analogues on Hmg2-GFP steady state levels in strains with elevated GGPP production. Same experiment in 4a, but with strains strongly expressing BTS 1 to increase GGPP and Hmg2-GFP degradation rate. Left panel strain expresses normally regulated Hmg2-GFP; right panel strain expresses unregulated TYFSA mutant of Hmg2-GFP.

We further explored the antagonistic action of GGSPP by examining its effects on GGPP-induced Hmg2 degradation *in vivo*. Because simultaneous addition of both GGPP and GGSPP could also have interactions on the unknown influx mechanism that appears to operate in yeast, we explored the effect of the inactive analogues on the endogenous GGPP degradation signal, which we have extensively characterized (20, 24). We first simply added each analogue to a strain with sufficient flux through the sterol pathway to produce the needed GGPP signal for Hmg2-GFP degradation. Specifically, we examined the effect of addition of inactive analogue on the Hmg2-GFP levels during a three hours incubation period. The effects of the analogues were small, but consistent with the *in vitro* effects of each: 2F-GGPP had no effect, while the GGSPP caused a small but reproducible increase in Hmg2-GFP steady-state (Fig 4A, left), implying that the added antagonist can block the degradation-stimulating effect of endogenous GGPP. Importantly, an identical experiment with the similarly degraded but unregulated TFYSA mutant of Hmg2-GFP showed no effect of the GGSPP antagonist on steady state levels, indicating that its effect was due to altering the response to GGPP signal, rather than effects on the HRD pathway itself (Fig 4A, right).

To further evaluate *in vivo* antagonism, we developed a yeast strain that constitutively generates high levels of endogenous GGPP, ensuring continuous strong signal, and thus as high a rate of regulated Hmg2-GFP degradation possible. Although the effect of GGPP was originally discovered by direct addition to living cultures, our studies confirmed that endogenous GGPP was responsible for regulating Hmg2 stability (20). Endogenous GGPP can be produced by several means, including through the action of the non-essential enzyme GGPP synthase, called Bts1 (35). In our previous work, we genetically manipulated the levels of the Bts1 by expressing it from the strong galactose-inducible GAL1 promoter. Capitalizing on this mode of GGPP generation, we made a yeast strain that constitutively expressed Bts1 from the similarly strong TDH3 promoter, to cause continuous endogenous production of high levels of GGPP. Expression of pTDH3-Bts1 decreased steady state Hmg2-GFP levels by about five-fold from wild-type strains (Fig 4B, left) and further addition of GGPP to culture media did not further decrease Hmg2-GFP (Fig 4B, right, orange curve), indicating that TDH3-Bts1 is producing maximally effective levels of GGPP. To confirm that the Bts1 was producing GGPP through the normal sterol pathway, we added the HMGR inhibitor lovastatin to block the normal production of the Bts1 substrates FPP and IPP. As expected, treatment with lovastatin increased Hmg2-GFP levels approximately six-fold (Fig 4B, right, blue curve). This constitutive high GGPP-producing strain further demonstrated the importance of GGPP in Hmg2 stability control, and allowed further testing its antagonism *in vivo*. Using the TDH3-Bts1 strain, we again tested the effects of direct addition of the GGPP analogues on Hmg2-GFP levels *in vivo* using flow cytometry. Consistent with the result from wild-type cells, 2F-GGPP did not change Hmg2-GFP levels, while the *in vitro* GGPP antagonist, GGSPP resulted in a nearly two-fold increase Hmg2-GFP steady state levels over the course of the incubation (Fig 4C, left). The expression of additional Bts1 had no effects on the *in vivo* levels of the non-responding mutant TFYSA. Again, neither analog had any effects on this unregulated protein (4C, right). Taken together, these results indicate that GGPP acts directly on the Hmg2 transmembrane domain, using a binding site with sufficient structural selectivity to show high potency, stringent structure activity, and specific antagonism both *in vivo* and *in vitro*.

*Testing GGPP as a ligand that promotes regulated misfolding*-We have previously proposed the idea that regulated Hmg2 degradation entails a programed or regulated change to a more unfolded form, thus enhancing the probability of entry into the HRD quality control pathway (22, 26, 36). This model is supported by the observed stabilization of rapidly degraded Hmg2 by the chemical chaperone glycerol. Addition of glycerol at concentrations required for chemical chaperoning (5-20%) causes rapid and reversible stabilization of Hmg2 that is undergoing *in vivo* degradation (24). Furthermore, the effect of glycerol is also observed in the in limited trypsinolysis assay (25). We had previously shown that the effects of high concentrations of farnesol on Hmg2 were reversed by glycerol, consistent with the other evidence that this lipid causes selective misfolding of Hmg2. Accordingly, we tested the ability of glycerol to antagonize the effects of the highly potent GGPP induced structural transition. First, we confirmed glycerol’s effects on Hmg2 levels *in vivo*.

Addition of glycerol at concentrations required for chemical chaperoning (typically 10-20%) directly to the culture medium increased Hmg2-GFP steady state levels (Fig 5a, left), and slowed the degradation rate as measured by cycloheximide chase (Fig 5A, right), as expected from our earlier studies. We then used glycerol to evaluate the role of misfolding in the action of GGPP. When cells were treated with maximal concentrations of GGPP and subjected to cycloheximide chase, Hmg2-GFP’s half-life drops from 1.5 hour to approximately 30 minutes. Co-addition of glycerol partially reversed the effects of added GGPP, increasing Hmg2-GFP’s half-life to over 1 hour (Fig 5B, left). We further tested the effect of glycerol using the *in vitro* proteolysis assay. We treated microsomes from cells expressing myc_L_-Hmg2-GFP with 20% glycerol, 27 μM GGPP, or both simultaneously. As expected from earlier work with less specific signals such as FOH (26), addition of glycerol decreased the effects of added GGPP, shifting the accessibility closer to that of untreated microsomes (Fig 5B, right).

**Figure 5.**
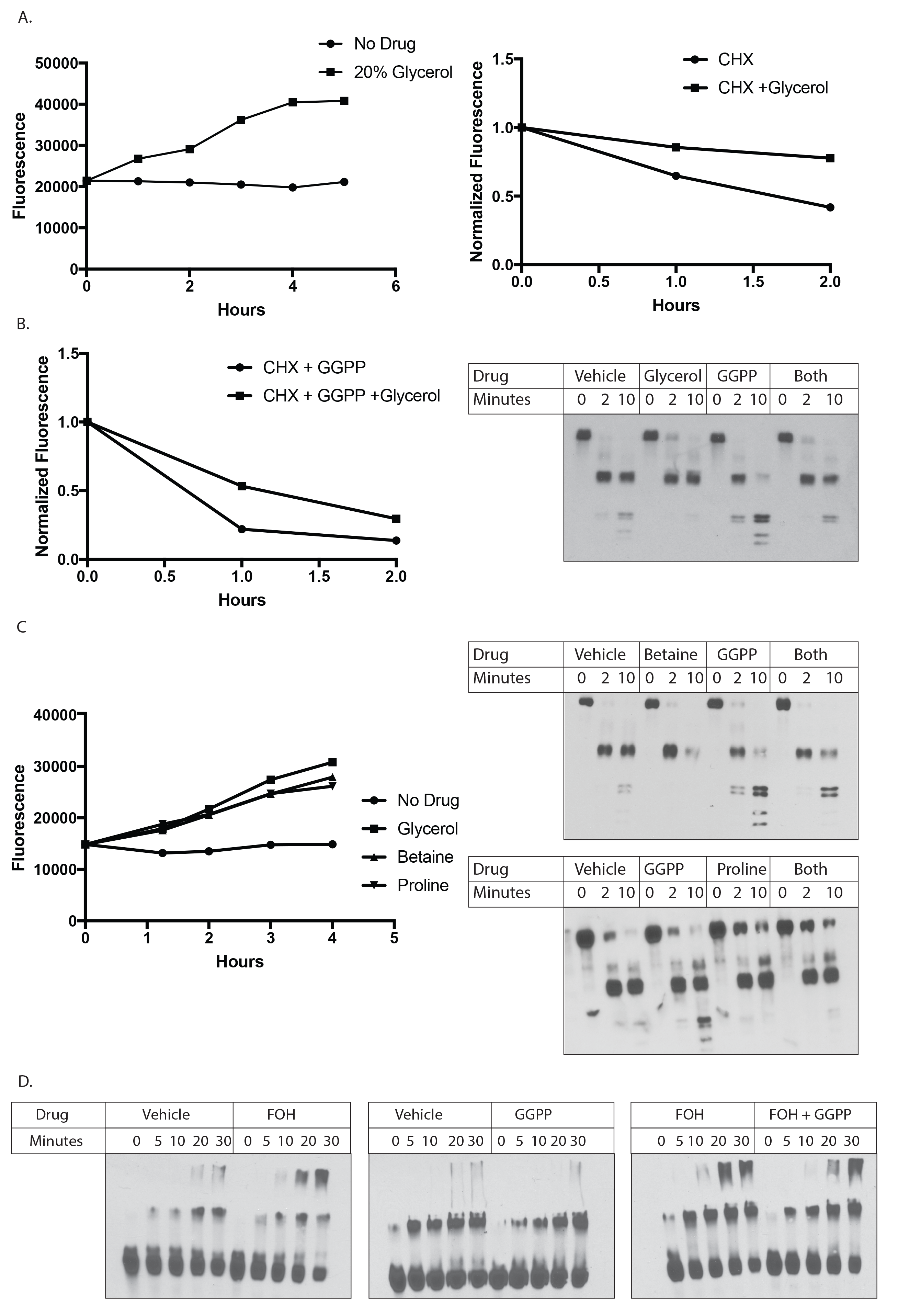
Effect of chemical chaperones on GGPP action *in vivo* and *in vitro*. (A) Effect of 20% glycerol on steady state levels of Hmg2-GFP (left) in living cells over the indicated times after addition, or the degradation rate of Hmg2-GFP (right) after addition of cycloheximide (CHX). In CHX chases, fluorescence is normalized to starting value at time of CHX addition. (B) Glycerol diminished the effect of added GGPP on Hmg2-GFP degradation as measured after CHX addition (left panel) or on Hmg2-GFP limited proteolysis due to trypisn (right panel). Glycerol was added to cells or microsomes immediately prior to the start of incubations. (C) Similar effect of two other chemical chaperones, betaine or proline, on Hmg2-GFP steady state in living cells (left panel) or Hmg2-GFP limited proteolysis (right panels), as indicated. (D) Effect of GGPP on Hmg2-GFP thermal denaturation, in the presence and absence of FOH. Note that GGPP does not cause enhanced thermal denaturation of Hmg2-GFP, but does mildly antagonize that caused by FOH.

These results with glycerol were consistent with GGPP causing remediable change in the folding state of Hmg2 both *in vivo* and *in vitro*, and occurred at concentrations consistent with its well-known action as a chemical chaperone. To confirm this misfolding model of GGPP, we next tested the effects of two entirely distinct chemical chaperones, proline and betaine. Each molecule similarly increased Hmg2-GFP levels *in vivo* (Fig 5D, left) and reversed the effect of GGPP *in vitro* (Fig 5D, right).

Another indicator of protein misfolding is increased susceptibility to thermal denaturation. We previously showed that treatment with high concentrations of FOH made Hmg2 more susceptible to denaturation, as indicated by the formation of low-mobility electrophoretic species during brief incubation at 70^°^ C (Shearer and Hampton 2005). We tested GGPP in our assay for thermal denaturation. Surprisingly, treating microsomes from a strain expressing Hmg2-GFP with GGPP at concentrations up to 20 μM did not lead to increased thermal denaturation or formation of low-mobility structures. Rather, treatment with GGPP actually slightly *decreased* thermal denaturation of Hmg2 compared to vehicle (Fig 5E, middle). Furthermore, when microsomes were treated with both GGPP and FOH, GGPP slightly antagonized the FOH-induced denaturation, providing additional evidence for ligand binding (Fig 5E, right). Thus, although GGPP causes an opening of the Hmg2 molecule similar to FOH, and this effect is reversed by chemical chaperone treatment, the degree of Hmg2 misfolding caused by the potent and physiological GGPP signal is clearly less extreme.

Since the GGPP-caused structural transition is reversible and antagonizable, we drew an analogy to allostery. By this model, GGPP binding to a specific site would alter the structure of Hmg2 to allow a more unfolded structure that is amenable to better recognition by the HRD machinery and reversible with chemical chaperones, but is not grossly misfolded. Although allosteric transitions are usually discussed with respect to enzyme kinetics or related protein functions, it is easily conceivable that a similar alteration in structure could render a substrate more or less susceptible to engagement of quality control machinery. Nearly all allosteric proteins are multimeric, and many require this structural feature for allostery to occur (37). We tested whether Hmg2 exists as a multimer using co-immunoprecipitation, modifying our method to analyze *in vivo* interactions of Hmg2 and other proteins (23, 38, 39). Specifically, we co-expressed Hmg2 tagged with GFP and Hmg2 with a myc tag in the linker domain in the same yeast strain. Co-expressing cells were subjected to non-detergent lysis and microsomes were prepared. Microsomes were then solubilized and Hmg2-GFP was immunoprecipitated. When both tagged constructs were co-expressed in the same strain, immunoprecipitation of Hmg2-GFP caused coprecipitation of 1myc-Hmg2, demonstrating that Hmg2 forms multimeric structures (Fig 6A). When only Hmg2-GFP or 1myc-Hmg2 was expressed in a strain, we were unable to detect the other tag in input lysates or immunoprecipitations (Fig 6A).

**Figure 6.**
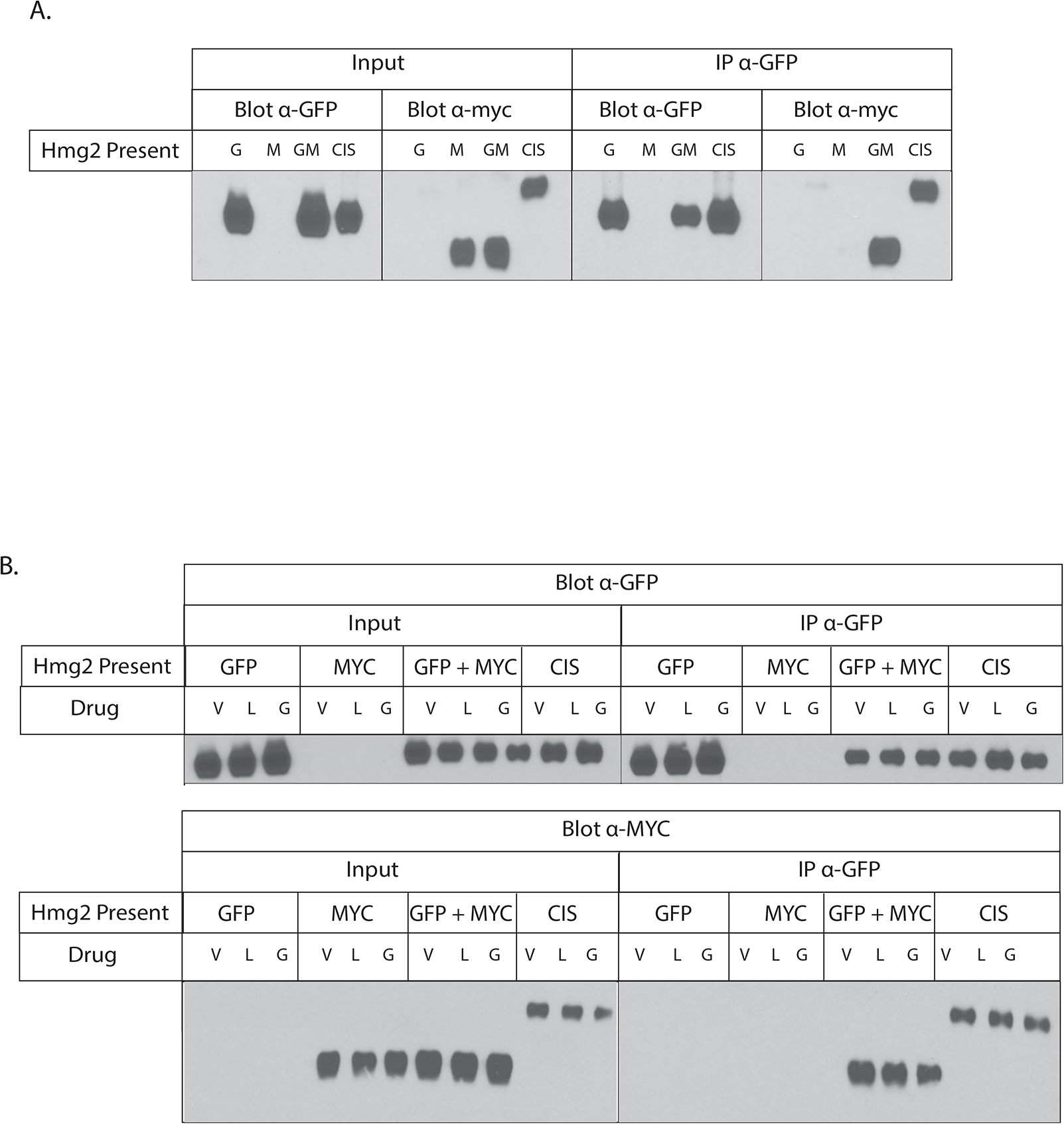
Hmg2 forms multimers *in vivo*. (A) Hmg2 transmembrane regions forms multimers. Co-immunoprecipitation of myc-tagged Hmg2 when co-expressed with Hmg2-GFP. Microsomes from strains expressing myc-Hmg2 (“M”), Hmg2-GFP (“G”), or both (“GM”) were solubilized with non-denaturing detergent, and subject to GPF immunoprecipitation (“IP α-GFP”), followed by immunoblotting for GPF or myc as indicated. Inputs of 10% total lysates are shown in the left group (“input”). “CIS” is a strain that expresses a single myc tagged Hmg2-GFP thus putting the myc tag and GFP in cis on the same protein, as a positive control. (B) (Lack) of effect of changing GGPP on co-precipitation of myc immunoblot. Experiments as in 6a, but with lovastatin pretreatment (L) as described, to lower GGPP, or added GGPP prior to and during immunoprecipitation to test if altering the regulator had any effect on Hmg2 self-association.

We asked if GGPP could affect Hmg2 multimerization. We repeated the co-immunoprecipitation experiments using cells treated with lovastatin to decrease GGPP levels *in* vivo, cells treated with added GGPP, or cells treated with only vehicle. In addition, lysis, microsome preparation, and co-immunoprecipiation from the GGPP-treated cells were all performed in the presence of added 22 μM GGPP, a concentration that maximally stimulates *in vivo* degradation and *in vitro* structural effects. In all three conditions, immunoprecipitating Hmg2-GFP pulled down the same amount of 1myc-Hmg2, suggesting that GGPP does not affect multimerization (Fig 6B).

Taken together, these data indicate that the regulation of Hmg2 entry into the widely used and conserved HRD quality control pathway occurs by a specific and reversible interaction with the naturally produced GGPP molecule. Furthermore, the structure-activity of this interaction is stringent, to the extent that closely related structures can antagonize the effects both *in vitro* and *in vivo*. A reasonable model for these data is that this represents a form of “folding allostery” by which the GGPP regulator causes a subtle structural transition to a more open or less folded state to promote physiological regulation by constitutive quality control.

## DISCUSSION

In these studies we sought to understand the GGPP-mediated regulation of Hmg2 ERAD. This included a detailed study of the structure-function features of GGPP, and the effects this biological regulator had on Hmg2 itself. The emerging model is that GGPP serves as a classic allosteric regulator that, instead of reversibly changing the parameters of enzyme action, instead causes reversible changes in folding state to bring about physiological regulation. The potential of this mode of regulation both for basic understanding and translational implementation are high.

Using flow cytometry, we tested a variety of isoprenoid molecules for their ability to induce Hmg2 degradation *in vivo*. We also used a limited proteolysis assay of Hmg2’s structure to directly examine the action of naturally occurring isoprenoids on Hmg2 structure. We found remarkable specificity for the 20-carbon isoprenoid GGPP both *in vivo* and *in vitro*. *In vivo*, GGPP was the only isoprenoid to induce Hmg2 degradation. *In vitro*, GGPP’s action was both highly potent and specific. The *in vitro* effect of GGPP on Hmg2 could be observed at concentrations as low as 12 nM, with a half-maximum concentration in the high nanomolar range, within an order of magnitude of the K_m_ of yeast enzymes which use GGPP, and thus consistent with its role as a physiological indicator of mevalonate pathway activity. Other isoprenoids tested required concentrations orders of magnitude higher to induce changes in Hmg2 folding in the limited proteolysis assay.

Consistent with its high potency, we found that the effect of GGPP showed extreme structural specificity. Two close analogs of GGPP, 2F-GGPP and GGSPP, despite being very similar biophysically, had no effect on Hmg2 structure at any concentration tested. In fact, one of the inactive molecules GGSPP was a GGPP antagonist: GGSPP interfered with GGPP’s effects on Hmg2 both *in vivo* and *in vitro*. Thus, the action of GGPP showed high potency, high specificity and was subject to inhibition by specific antagonist analog. Taken together, these observations suggest that GGPP controls Hmg2 ERAD by binding to a specific site on the Hmg2 transmembrane region, much like an allosteric regulator of an enzyme.

We also examined the nature of Hmg2’s response to GGPP. Because Hmg2 undergoes regulated entry into the HRD quality control pathway, our early studies examined whether Hmg2 undergoes regulated misfolding to make it a better HRD substrate. Consistent with this model, we showed that the chemical chaperone glycerol causes striking elevation of Hmg2 stability *in vivo* and drastically slows the rate of Hmg2 limited proteolysis (25). Those early studies showed that the 15 carbon isoprenoid molecule farnesol (FOH) caused Hmg2 to become less folded, and this effect of FOH was not observed with mutants of Hmg2 that do not undergo regulated degradation *in vivo* (26). The *in vitro* effect of FOH was antagonized by chemical chaperones, indicating that FOH causes Hmg2 misfolding (26). At the time of those studies, we did not know about GGPP, and found FOH’s specific but fairly impotent effects by direct tests *in vitro*.

Accordingly, in these current studies we explored if the more potent and biologically active GGPP similarly caused programmed misfolding. Indeed, glycerol reversed the effects of GGPP both *in vivo* and *in vitro*. We also tested two other distinct chemical chaperones—proline, betaine. These also prevent Hmg2 *in vitro* misfolding and *in vivo* degradation upon GGPP treatment. The generality of these chemical chaperones’ effects suggested that Hmg2’s entry into a quality control is mediated by regulated misfolding of Hmg2 in response to GGPP.

In those early studies exploring the effects of FOH on Hmg2, we also used thermal denaturation as a gauge of *in vitro* Hmg2 misfolding. Incubation of microsomes at 70 C induced aggregation of Hmg2 into a high molecular weight, denatured form that remains in the stacking gel of an SDS-PAGE separation, allowing straightforward assessment of time-dependent thermal denaturation by immunoblotting (26). We showed that treatment of microsomes with high micromolar concentrations of farnesol increased the rate and extent of Hmg2 thermal denaturation, while mutants of Hmg2 that do not respond to FOH in the proteolysis assay also did not show effects of FOH on thermal denaturation. In contrast, GGPP did not affect Hmg2 thermal denaturation, and may in fact have a slight protective effect. GGPP treatment also partially antagonized the thermal denaturation caused by farnesol. These combined results suggest that GGPP caused a subtler from of misfolding that is still remediable by chemical chaperones but not prone to enhance wholesale aggregation. In other words, GGPP action is a misfolding “sweet-spot”, sufficient to enhance selective degradation by the HRD machinery, but not the stress-inducing and health-compromising effects of wholesale misfolding or aggregation.

Combined, these features lead us to a model of “folding state allostery”, in which GGPP plays the role of an allosteric ligand. Upon interacting with GGPP, Hmg2 undergoes a conformational change to a partially misfolded state that renders it more susceptible to HRD degradation. GGPP meets the criteria for ligand-like behavior—its action is specific, potent, reversible, antagonizable, and occurs at physiologically relevant concentrations. Although usually allosteric regulators are view as “agonists” of a particular structural response, antagonizing ligands are often observable in classical enzyme allostery, and in fact can be part of the bone fide physiological control of enzyme activity in the cell. For example, AMP activates the kinase AMPK, but ATP competes to block this activation (40, 41).

GGPP-mediated Hmg2 misfolding is sufficient to gain entry to the ERAD quality control pathway, and can be reversed by treatment with several different chemical chaperones both *in vitro* and *in vivo*. However, this misfolding is not so severe as to make Hmg2 more thermally unstable and prone to aggregation. We thus propose, with admitted linguistic license, to call this structural effect “mallostery”-a portmanteau of the preface “mal” for misfolded or poorly structured- and allostery for the nature of this regulated and physiologically useful folding transition. While traditionally allostery has been viewed as a rigid phenomenon of highly ordered proteins, advances in structural methods have allowed for a more inclusive view. In recent years it has been more widely recognized that allosteric regulation occurs across the whole spectrum of order in proteins, including intrinsically disordered proteins. Allostery can capitalize on disorder and misfolding, with allosteric proteins undergoing disorder switching, local unfolding, or becoming partially disordered upon posttranslational modification (42, 43).

Because most allosteric proteins are multimeric, we tested whether Hmg2 forms multimeric structures, and found that indeed it does, but making use of co-expressed, fully regulated versions of Hmg2 with distinct epitopes. Furthermore, we found no evidence for alteration in multimerization caused by addition of even saturating concentrations of GGPP in the co-immunoprecipitation experiments. This also speaks to the idea of GGPP causing a more subtle change in structural state: full dissocation of a monomer caused by a ligand could certainly enhance recognition by the HRD pathway. However, again, it appears that the GGPP induced effects do not take things this far down the road to structural squalor. We picture the multimeric structure as allowing a subtle alteration of folding state that can be reversed upon removal of the GGPP ligand, thus allowing quality control regulation with minimal aggregation or denaturation.

A longstanding open question about Hmg2 has been whether its regulated degradation is due to binding of a ligand or rather due to a more global biophysical processes—for example, perturbation of the ER membrane by isoprenoid regulators. In this work, we find that GGPP causes Hmg2’s structural transition at nanomolar concentrations, far below the concentrations that would be expected to alter phospholipid bilayers properties. Furthermore, highly similar molecules were unable to affect Hmg2 at concentrations hundreds of times higher: While < 100 nM GGPP had clear effects on Hmg2 structure, 40 uM 2F-GGPP, had no discernable effect. In the same vein, our prior work has found a similarly striking degree of specificity for Hmg2 itself. Single point mutations within Hmg2 render it stable *in vivo* and unable to respond to GGPP *in vitro*. The combination of stringent sequence specificity for Hmg2, structural specificity for ligand, and the high potency of GGPP make it unlikely that Hmg2 misfolding and degradation the result of any general biophysical perturbation of the ER membrane proteome.

Another possible explanation for GGPP’s action is that, by binding to Hmg2, it presents a hydrophobic patch which the quality control machinery detects as the exposed core of misfolded protein. Such so-called “greasy patches” are the basis of a strategy for artificially engineering the degradation of target proteins (44, 45). This model is at odds with two observations. One, GGPP induces not only engagement with the UPS machinery, but a structural change *in vitro*. Two, very similar (and with an identical hydrophobic tail) molecules did not cause any effects on Hmg2 *in vitro* or *in vivo*. Furthermore, GGSPP, which is nearly identical to GGPP, antagonizes GGPP’s effects. This implies that GGSPP can bind at the same location as GGPP, but despite this it is unable to induce the structural transition or degradation. Were the hydrophobic end of GGPP the key to its action, one would not expect such a similar molecule to behave in the opposite manner.

This model leaves several open questions. We find that GGPP’s effects on Hmg2 are specific, potent, rapid, and reversible, but does GGPP bind directly to Hmg2, and if so, where? Are other ER proteins required for Hmg2 misfolding? It seems unlikely that there are unknown stoichiometric binding partners required for Hmg2 misfolding, as Hmg2 is overexpressed in our *in vitro* experiments, but the possibility remains. Furthermore, we find that Hmg2 can be co-immunoprecipitated with differently tagged Hmg2 constructs expressed in the same cell. If Hmg2 is a multimer, does the multimerized state influence this mode of regulation? Do members of the complex influence each other in undergoing the conformational change to the misfolded state, as in classical models of allostery, or are individual Hmg2s independent?

Thirty percent of the US population suffers from dyslipidemia; more have dyslipidemia controlled by pharmaceutical treatment (46). As the rate controlling step of the sterol pathway, HMGR is a key intervention point in metabolic disease; over 25% of adults in the US take cholesterol lowering medications, and over 20% statin drugs which target this protein (47, 48). Mammalian HMGR levels rise after statin treatment due to both increased transcription of sterol genes and stabilization of HMGR itself when sterol levels are low (49). Key components of HMGR regulated degradation are conserved in mammals, including induction by a 20 carbon isoprenoid and ubiquitination by ER E3 ligases, including gp78 which is a Hrd1 homologue (50–53). Furthermore, when the mammalian HMGR and its ancillary regulatory proteins are expressed in insect cells, endogenous Hrd1, the same E3 ligase as in yeast, mediates sterol-regulated HMGR degradation (54, 55). These extensive similarities in the system highlight the importance of a deeper understanding of the dynamics underlying HMGR’s regulated quality control degradation. A greater understanding the underpinnings of HMGR stability may open up avenues for better targeting the pathway in human patients.

This phenomenon of ligand-programmed misfolding raises questions about more general pharmacological applications. Proteins without active sites for inhibitors to engage can be difficult to target pharmacologically, and have in fact been referred to as the “undruggable proteome” (56). The UPS system has already been tapped as a tool for pharmacological targeting of these undruggable protein through regulated degradation. Two main strategies have emerged so far: targeting proteins directly to specific E3 ligases, such as VHL (57), and targeting proteins with ligands fused to a long hydrophobic molecule, or “greasy patch,” to mimic a misfolded protein (45). Directing proteins specifically to quality control by cleaver discovery of mallosteric regulators that cause selective unfolding may offer another approach for targeting the undruggable proteome, and one that nature has clearly already discovered during evolution.

## EXPERIMENTAL PROCEDURES

Reagents-Geranylgeranyl pyrophosphate (GGPP), geranylgeraniol (GGOH), farnesyl pyrophosphate (FPP), farnesol (FOH), geranyl pyrophosphate (GPP), isopentenyl pyrophosphate (IPP), and cycloheximide were purchased from Sigma-Aldrich. Lovastatin was a gift from Merck & Co (Rahway NJ). S-thiolo-GGPP (GGSPP) and 2-fluoro-GGPP (2F-GGPP) were gifts from Reuben Peters (Iowa State University) and Philipp Zerbe (University of California Davis). Anti-myc 9E10 supernatant was produced from cells (CRL 1729, American Type Culture Collection) grown in RPMI1640 medium (GIBCO BRL) with 10% fetal calf serum. Living colors mouse anti-GFP monoclonal antibody was purchased from Clontech. Polyclonal rabbit anti-GFP antibody was a gift from C. Zucker (University of California San Diego). HRP-conjugated goat antimouse antibody was purchased from Jackson ImmunoResearch. Protein A Sepharose beads were purchased from Amersham Biosciences.

*Yeast strains and plasmids*-Yeast strains (Table 1) and plasmids (Table 2) were constructed by standard techniques. The integrating Bts1 overexpression construct, plasmid pRH2657 was made by replacing the SpeI-SmaI fragment of pRH2654 with the Bts1 coding region amplified from pRH2477.

Yeast strains were isogenic and derived from the S288C background. Yeast strains were grown in minimal media (Diffco yeast nitrogen base supplemented with necessary amino and nucleic acids) with 2% glucose or rich media (YPD). Strains were grown at 30°C with aeration. Lumenally myc-tagged Hmg2-GFP constructs, were introduced by integration of plasmid cut with *StuI* at the *ura3-52* locus. The Bts1 overexpression construct was introduced by integration of plasmid cut with *PpuMI* at the *leu2Δ* promoter.

*Flow* cytometry-Flow cytometry was performed as described previously (24, 58). Briefly, yeast strains were grown in minimal media into early log phase (OD_600_<0.2) and incubated with the indicated isoprenoid molecules (naturally occurring isoprenoid concentrations as indicated; 44 μM GGSPP and 2F-GGPP unless indicated otherwise), drugs (25 μg/mL lovastatin and 50 ug/mL cycloheximide), or equal volumes of vehicle (for isoprenoid pyrophosphate molecules, 7 parts methanol to 3 parts 10 mM ammonium bicarbonate; for lovastatin, 1 part ethanol to 3 parts Tris Base pH 8; and for cyclohexmide, GGOH, and FOH, DMSO) for the times indicated. Individual cell fluorescence for 10,000 cells was measured using a BD Accuri C6 flow cytometer (BD Biosciences). Data were analyzed using FlowJo software (FlowJo, LLC).

*Microsome Preparation*-Microsomes were prepared as described previously (25). Yeast strains were grown to mid-log phase in YPD. 10 OD equivalents were resuspended in 240 μL lysis buffer (0.24 M sorbitol, 1 mM EDTA, 20 mM KH2PO4/K2HPO4, pH 7.5) with PIs (2 mM phenylmethylsulfonyl fluoride and 142 mM tosylphenylalanyl chloromethyl ketone). Acid-washed glass beads were added to the meniscus and cells were lysed at 4°C on a multi-vortexer for six 1-minute intervals with 1 minute on ice in between. Lysates were cleared by centrifugation in 5s pulses until no pellet was apparent. Microsomes were pelleted from cleared lysates by centrifugation at 14,000 x g for five minutes, washed once in XL buffer (1.2 M sorbitol, 5 mM EDTA, 0.1 M KH2PO4/K2HPO4, pH 7.5), and resuspended in XL buffer.

*Limited proteolysis assay*-Microsomes were subjected to limited proteolysis as described previously (25). Briefly, resuspended microsomes were treated with the indicated isoprenoid molecules or equal volumes of vehicle and then incubated with trypsin at a final concentration of 15 μg/mL at 30°C. Reactions were quenched at the times indicated with an equal volume of 2x urea sample buffer (USB; 8M urea, 4% SDS, 1mM DTT, 125 mM Tris base, pH 6.8). Samples were resolved by 14% SDS-PAGE, transferred to nitrocellulose in 15% methanol, and blotted with 9E10 anti-myc antibody.

*Thermal denaturation* assay-The thermal denaturation assay was performed as described previously (26). Briefly, resuspended microsomes were treated with the indicated isoprenoid molecules or equal volumes of vehicle and transferred to PCR tubes. Samples were placed in a thermocycler (Eppendorf Mastercycler Pro) preheated at 70°C and incubated at 70°C for the indicated times. Samples were held on ice for two minutes prior to addition of equal volumes of 2x USB. Samples were resolved by 14% SDS-PAGE, transferred to nitrocellulose in 10% methanol, and blotted with 9E10 anti-myc antibody.

*Microsome preparation for co-immunoprecipitation-Microsomes* were prepared for co-immunoprecipitation as described previously (39). Yeast strains were grown to midlog phase in YPD. 10 OD equivalents were resuspended in 240 μL lysis buffer (0.24 M sorbitol, 1 mM EDTA, 20 mM KH2PO4/K2HPO4, pH 7.5) with protease inhibitors (2 mM phenylmethylsulfonyl fluoride, 100 mM leupeptin hemisulfate, 76 mM pepstatin A, and 142 mM tosylphenylalanyl chloromethyl ketone). Acid-washed glass beads were added to the meniscus and cells were lysed at 4°C on a multi-vortexer for six 1-minute intervals with 1 minute on ice in between. Lysates were pelleted by centrifugation in 5-second pulses until no pellet was apparent, with the supernatant moved to a clean tube each time. Microsomes were pelleted from cleared lysates by centrifugation at 14,000 x g for five minutes, washed once in IP buffer without detergent (500 mM NaCl, 50 mM Tris base, pH7.5), and resuspended in IP buffer with detergent (IPB; 500 mM NaCl, 50 mM Tris base, 1.5% Tween-20, pH7.5) and protease inhibitors.

*Co-immunoprecipitation-Microsomes* in IBP were incubated at 4°C for 1 hour with rocking. Microsomes were pipetted up and down repeatedly and then solutions were cleared by centrifugation at 14,000 x g for 15 minutes. Supernatants were incubated with 15 μL polyclonal rabbit anti-GFP antibody overnight at 4°C with rocking. After overnight incubation, 100 μL of 50% protein-A sepharose bead slurry swelled in IP buffer without detergent were added. Samples were incubated at 4°C for two hours with rocking. Beads were then pelleted for 30 seconds at low speed and 1 minute by gravity and washed twice with IPB and once with IP wash buffer (100 mM NaCl, 10 mM Tris base, pH7.5. Beads were aspirated to dryness and resuspended in 2x USB. Samples were resolved by electrophoresis on 14% polyacrylamide gels, transferred to nitrocellulose in 12% methanol buffer, and immunoblotted with anti-GFP or anti-myc antibody as indicated.

## ACKNOWLEDGEMENTS

We thank Reuben Peters (Iowa State University) and Philipp Zerbe (University of California Davis) for providing reagents. These studies were supported by NIH grant 5R37DK051996-18 to R.Y.H. M.A.W. was supported in part by NIH CMG Training Grant 5T32GM007240-35.

